# Deep learning-enhanced morphological profiling predicts cell fate dynamics in real-time in hPSCs

**DOI:** 10.1101/2021.07.31.454574

**Authors:** Edward Ren, Sungmin Kim, Saad Mohamad, Samuel F. Huguet, Yulin Shi, Andrew R. Cohen, Eugenia Piddini, Rafael Carazo Salas

**Affiliations:** School of Cellular & Molecular Medicine, University of Bristol, BS8 1TD Bristol, UK; Department of Electrical and Computer Engineering, Drexel University, 3120-40 Market Street, Suite 313, Philadelphia, PA 19104, USA.

**Keywords:** human Pluripotent Stem Cells, Single-cell microscopy phenomics, Morphological profiling, Live multi-colour microscopy, Multiday imaging, High-content microscopy, Computational image analysis, Deep Learning, Neural networks, Cell fate dynamics prediction.

## Abstract

Predicting how stem cells become patterned and differentiated into target tissues is key for optimising human tissue design. Here, we established DEEP-MAP - for deep learning-enhanced morphological profiling - an approach that integrates single-cell, multi-day, multi-colour microscopy phenomics with deep learning and allows to robustly map and predict cell fate dynamics in real-time without a need for cell state-specific reporters. Using human pluripotent stem cells (hPSCs) engineered to co-express the histone H2B and two-colour FUCCI cell cycle reporters, we used DEEP-MAP to capture hundreds of morphological- and proliferation-associated features for hundreds of thousands of cells and used this information to map and predict spatiotemporally single-cell fate dynamics across germ layer cell fates. We show that DEEP-MAP predicts fate changes as early or earlier than transcription factor-based fate reporters, reveals the timing and existence of intermediate cell fates invisible to fixed-cell technologies, and identifies proliferative properties predictive of cell fate transitions. DEEP-MAP provides a versatile, universal strategy to map tissue evolution and organisation across many developmental and tissue engineering contexts.

## INTRODUCTION

How to define cellular states and state transitions remains a fundamental question in stem cell biology. In recent years, approaches that can quantify properties of stem cells and their differentiated derivatives have become broadly recognised for their pivotal roles in predictable and robust synthetic tissue design (Brassard and Lutolf, 2019; Del Sol et al., 2017; Prochazka et al., 2017). Although RNAseq- and DNA barcoding-based single-cell technologies are used extensively for cell state mapping and lineage analysis (Cahan et al., 2014; Kinney et al., 2019; Rackham et al., 2016; VanHorn and Morris, 2021), these have critical limitations: cells are killed and their characteristics are measured only at that point in time. Hence, historical cell state information that might be key to fate evolution – e.g. transient cell states that disappear before the endpoint observation, dying cells at the time of observation, cell tissue context and density prior fate acquisition – is lost. Furthermore, for single-cell technologies time is an implicit variable that can only be inferred mathematically. This is important, because the true dynamical and temporal variables (individual and collective/reciprocal cell movements, the actual duration and longevity of cell states, and how cell heterogeneity changes and co-evolves with fate) is lost as well (Villoutreix, 2021).

In contrast, continuous live cell imaging provides a non-interventional interrogation of cell state dynamics, including historical information of cell states and cell state transitions, in real-time (Chessel and Carazo Salas, 2019). ‘Live’ tracking of cell states has so far relied on using fluorescently-tagged transcription factors to reveal cell fate/state (Etzrodt and Schroeder, 2017; Filipczyk et al., 2015; Strebinger et al., 2019; Wolff et al., 2018), although newer strategies using indirect reporters of transcription factor status are emerging (Kim et al., 2021). Crucially, each differentiation or other experimental scenario requires the establishment of a tailor-made reporter. This limitation becomes prohibitive when aiming to study cell differentiation into multiple target fates, as the possible reporters that can be co-expressed and imaged ‘live’ simultaneously in cells are relatively few (Kim et al., 2021). Developing technologies to quantitatively monitor cell state dynamics in real-time and without the need for specialised cell state reporters would overcome these limitations and provide a powerful solution to study and predict cell state transitions.

In the past 15 years, high-dimensional, image-based morphological profiling enhanced by machine learning has been used successfully by many groups including ours in a variety of model systems to systematically identify and characterise genes/gene-network states (Chong et al., 2015; Collinet et al., 2010; Fuchs et al., 2010; Graml et al., 2014; Neumann et al., 2010), to identify and enhance signalling networks (Bakal et al., 2007; Evans et al., 2013; Horn et al., 2011), to characterise small molecule treatments (Bray et al., 2017; Bray et al., 2016) and to infer context-dependent gene functions (Sailem et al., 2020). To date, however, machine learning-enhanced ‘live’ morphological profiling has not been used to identify and predict cell fate transitions in real-time.

Here, we established DEEP-MAP, an approach that combines optimised multi-day, multi-colour microscopy phenomics with deep neural networks, and allows to harness high-dimensional morphological profiling information obtained from hundreds of thousands of ‘live’ cells, to map and predict cell fate dynamics in real-time. We applied DEEP-MAP to human pluripotent stem cells (hPSCs) co-expressing broadly used cell proliferation and cell cycle reporters and asked whether phenotypic profiling alongside proliferation information can be used to identify cell states and monitor multi-fate dynamics. We show that based only on morphological and proliferative features DEEP-MAP can reliably map and spatiotemporally predict, at a single-cell level, cells’ fate and fate dynamics during differentiation into the three basic germ layer fates, without a need for customised cell state reporters specific to the cell fates and lineages being monitored. We show that DEEP-MAP predicts fate changes with high temporal sensitivity - comparable to what is currently technically possible using fluorescently-labelled transcription factor reporters. Moreover, we show that DEEP-MAP can reveal the timing of phenotypic transitions associated with cell fate conversions and the existence of cell fate intermediates, and be used to identify proliferative properties predictive of cell fate transitions.

## RESULTS

### Establishing a multi-day, multi-colour microscopy phenomics pipeline for dynamical phenotyping of hPSCs in real-time and at single-cell resolution

To monitor hPSC morphology and proliferation dynamics at single-cell resolution, we generated a CRISPR knock-in three-colour hPSC line co-expressing a fluorescently-tagged histone H2B reporter, H2B-miRFP670 (Kim et al., 2021; Shcherbakova et al., 2016) (far red fluorescence emitting), and the two-colour FUCCI cell cycle reporter (Pauklin and Vallier, 2013; Sakaue-Sawano et al., 2008) (red/green fluorescence emitting) (Fig. 1A) (see STAR Methods for detailed descriptions from here on). When considering choice of reporters, we took into account that (1) hPSCs form very compact colonies; therefore fluorescent nuclear reporters would give the best shot at unequivocal single-cell identification and morphological profiling (2) FUCCI and H2B are robust cell proliferation reporters that together enable quantitative monitoring of detailed aspects of cell proliferation, cell cycle progression, mitosis and cell death and (3) the H2B signal specifically never disappears from cells, allowing uninterrupted cell visualisation. We then set out to establish a multi-colour, time-lapse high-content microscopy pipeline enabling us to image cells continuously over time, in a way that would be compatible with daily media change, high-temporal resolution imaging (to allow automated cell detection and tracking) and multi-day imaging, required to observe fate transitions occurring on a time scale of days as cells continue to proliferate normally (Fig. 1B).

**Figure 1.**
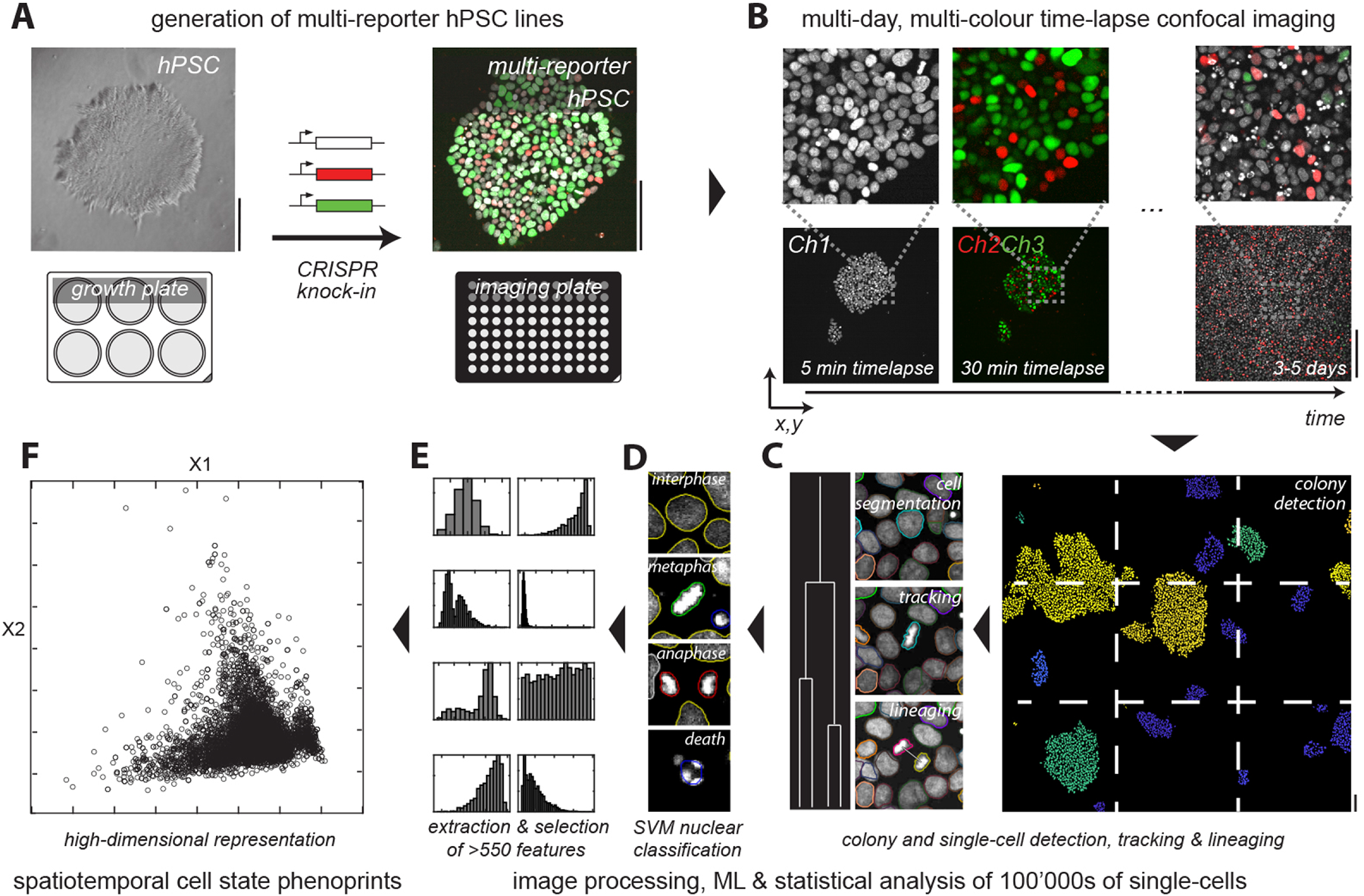
Establishing a multiday high-content microscopy pipeline to quantitively profile ‘live’ hPSC dynamics in real-time and at single-cell resolution. **A.** We used CRISPR knock-in to generate a three-colour, multi-reporter cell line co-expressing FUCCI and fluorescently-tagged H2B, enabling comprehensive monitoring of hPSCs morphology and proliferation during pluripotency and early differentiation. Scalebars: 100 μm. **B.** Multi-colour confocal imaging over up to five days was achieved by differential time-lapse sampling of fluorescence channels (fluorescently-tagged H2B every 5 min, FUCCI every 30 min), allowing us to capture both short-time information needed for cell detection and tracking as well as longer-time proliferative changes observed during fate transitions while keeping cells healthy under the microscope. Scalebar: 200 μm. **C, D** and **E.** Images from neighbouring fields, digitally stitched into larger images containing multiple hPSC colonies, were used to computationally detect and track colonies over time (**C**, right), where possible, detect and track cells over time (**C**, left), classify cells into interphase, metaphase, anaphase or dead cells by machine learning (**D**), and extract >550 different morphological, intensity and texture features across the multi-channel signals on a single-cell basis (**E**). **F**. This pipeline allowed us to obtain high-dimensional morphological and proliferation phenoprints for hundreds of thousands of cell datapoints for each experimental condition analysed.

Frequent and extended time-lapse imaging is highly phototoxic (Loeffler and Schroeder, 2019; Piltti et al., 2018; Schroeder, 2011), particularly for stem cells, prohibiting imaging many reporters with high temporal frequency. To overcome this limitation, we developed an optimised imaging modality by which we imaged the H2B-miRFP670 signal every 5 minutes and the FUCCI signal only every 30 minutes. This scheme allowed us to achieve multi-day imaging of healthily proliferating cells for at least 3-5 days or more, limited only by cell confluence. In this manner, we captured multi-colour time-lapse images of cell populations across multiple image fields over time, and then used a customised image analysis pipeline to (A) digitally stitch neighbouring fields into larger images containing multiple hPSC colonies 1. (B) detect all cells in all colonies through time (using nuclei as proxies; from here onwards cells/nuclei are used interchangeably) (C) where possible, track individual cells through time, and (D) detect and track all colonies through time (Fig. 1C). Typically, time series captured in this way consisted of 800-1000 images (timesteps), with each imaging field leading to >100,000 detected nuclei data points throughout the entire multi-day imaging sequence. After data capture, we used a trained multi-class Support Vector Machine (SVM) to classify and assign probabilities by machine learning to different types of nuclear events (interphase, metaphase, anaphase, cell death) (Fig. 1D), and then extracted, in addition to the SVM-derived features (i.e. probability assignments), >550 different morphological, intensity and texture features across the multi-channel signals on a single-cell basis (Fig. 1E). Ultimately, this analysis yielded >550-dimensional phenoprints for hundreds of thousands of cells for each experimental condition analysed, capturing the morphological and spatiotemporal phenotype of that condition (Fig. 1F).

### Live morphological profiling of hPSCs during pluripotency and early germ layer differentiation

Next, we used this strategy to dynamically phenotype the evolution of hPSCs, either while maintaining pluripotency or when triggered to undergo early directed differentiation. Given that hPSCs have the capacity to differentiate into all three basic germ layers, we chose to trigger cells to undergo directed differentiation into primitive neural stem cells (NSCs), cardiac mesoderm-induced cells, and definitive endoderm, mimicking early ectoderm (EC), mesoderm (ME) and endoderm (EN), respectively (Fig. S1; hereafter referred to as EC, ME and EN for shorthand). In our conditions, pluripotent hPSCs co-expressing H2B-miRFP670 and FUCCI formed colonies that grew exponentially and divided with a typical cell doubling time of 18 hours, in agreement with an average cell cycle duration of ∼15 hours (Pauklin and Vallier, 2013), with most cells in S/G2/M (FUCCI-green) phases of the cell cycle (Fig. S1A and Movie S1). Similarly, hPSCs triggered to initiate EC differentiation initially grew exponentially with a typical doubling rate of 18 hours in the first 48 hours, after which they visibly slowed their doubling time while gradually increasing the proportion of cells in G1 (FUCCI-red) (Fig. S1B and Movie S2), coincident with differentiation onset. By contrast, cells triggered to differentiate into ME (Fig. S1C and Movie S3) and EN (Fig. S1D and Movie S4) changed their appearance after 24h of imaging, becoming more spread out and altering their nuclear shape and cell cycle reporter characteristics, especially in EN triggered cells (Figs. S1C-S1D).

To compare the phenotypic evolution of the four different cell populations quantitatively, we sought to map the high-dimensional phenoprints of the four cell populations. We found that linear dimensionality reduction by principal component analysis (PCA, (Pearson, 1901) did not separate the four populations well. This outcome suggested that population variance in feature space was overall comparable and overlapping (Fig. 2A), possibly due to the low signal-to-noise (SNR) ratio imposed by the low exposure times necessary in our time-lapse imaging to keep cells healthy (Weigert et al., 2018). Similarly, when we used non-linear embeddings commonly used for single-cell transcriptomics analysis (Kobak and Berens, 2019), t-distributed stochastic neighbour embedding (tSNE (van der Maaten and Hinton, 2008)) and Uniform Manifold Approximation and Projection (UMAP, (McInnes et al., 2018), we found that (with the exception a subset of EN triggered cells) the four cell populations largely overlapped in those embedding spaces (Figs. 2B-2C).

**Figure 2.**
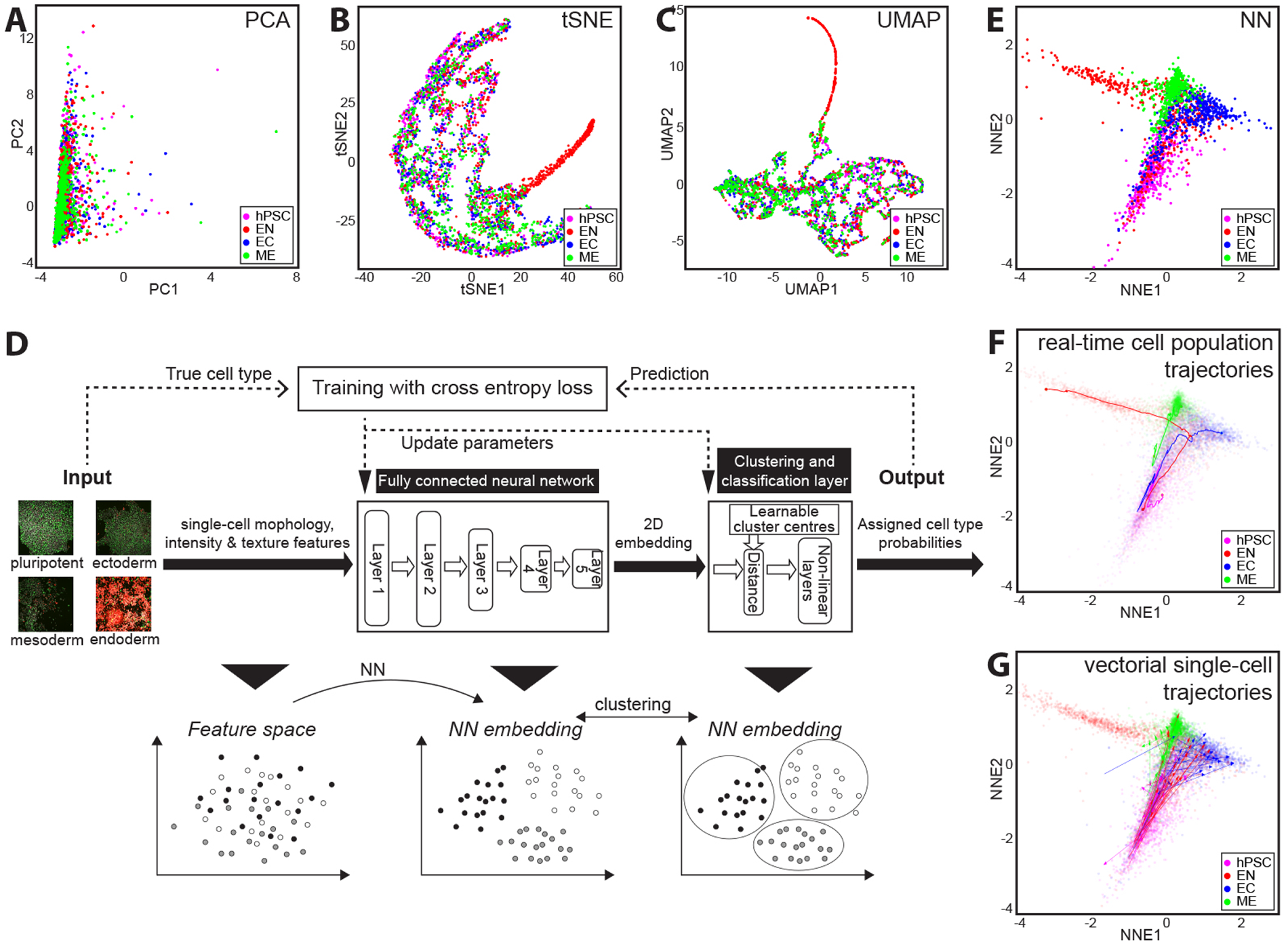
Mapping and clustering cell fate dynamics by deep learning. **A**, **B** and **C**. Mapping high-dimensional phenoprints of the four cell populations analysed (hPSC, EC, ME and EN) by PCA (**A**), tSNE (**B**), and UMAP (**C**) did not yield good clustering or population separation, likely due to the low SNR of the time-lapse imaging derived data. **D**. Flowchart outlining the design of a neural network architecture for predicting and generating a spatial mapping (embedding) of low SNR morphological profiling data. **E**. Neural networks allow clear mapping and separation of cell populations based on their morphological and proliferative phenoprints. **F**. Mapping the dynamic evolution and phenotypic diversity of the four cell populations in real-time. Solid lines: population trajectories; magenta/red/blue/green: hPSC/EN/EC/ME cell populations, correspondingly. Real-time trajectories are shown as solid lines overlaid on the neural network embedding in **E** (made partly transparent for visualisation purposes). **G**. Vectorial trajectories of single-cells tracked early in differentiation along the different lineages. Arrows: Vectorial trajectories; magenta/red/blue/green: hPSC/EN/EC/ME cell populations, correspondingly. Single-cell trajectories are shown as solid arrows overlaid on the neural network embedding in **E** (made partly transparent for visualisation purposes).

### Using neural networks to map phenotypic diversity

To overcome these methodological limitations, we developed a supervised neural network (NN) embedding model that first uses the >550-dimensional feature datasets to learn to predict and spatially embed different cell state classes as separate as possible on a plane knowing their class labels, and can then be used subsequently to map new data points based on their feature phenoprints without knowledge of the class labels (Fig. 2D). We trained the model with four input classes: ‘pluripotent’, ‘ectoderm’, ‘mesoderm’ and ‘endoderm’. ‘Pluripotent’ consisted of 2,675 early pluripotent datapoints (t=0h-6h, i.e. 0 days, when colonies are small) and 23,492 late pluripotent datapoints (t=60h40min-66h40min, i.e. 2.5 days, when colonies are visually large and dense). Importantly, the ‘pluripotent’ training class contained both early and late pluripotent cell datapoints to eliminate cell density as a potential confounding factor. ‘Ectoderm’ consisted of 23,659 late EC datapoints (t=77h20min-83h20min i.e. 3.5 days, when cells acquire SOX1+, denoting early EC fate (Kim et al., 2021)). ‘Mesoderm’ consisted of 15,278 late ME datapoints (t=77h20min-83h20min i.e. 3.5 days, when cells have been reported to exit the T+ state en route to becoming early cardiac mesoderm fate (Rao et al., 2016)). ‘Endoderm’ consisted of 1’296 late EN datapoints (t=60h40min-66h40min i.e. 2.5 days, when cells have been shown to become SOX17+ denoting early EN fate (Ogawa et al., 2013; Teo et al., 2011)). This approach proved to be pivotal for clearly separating the different cell states (Fig. 2E). Crucially, we were able to reproducibly do so across different datasets (Fig. S2A-S2I), showing that the NN model is both predictive and robust.

To visualise the dynamic evolution of the different cell populations and exploit the real-time nature of our data, we then computed the average coordinates of each cell population at every time point for the entire 3.5 days to generate their real-time phenotypic trajectories (Fig. 2F). As expected, at early timepoints (0-1 days), all four cell populations were close to each other and located around the pluripotent embedding area, corresponding to the fact that they were all phenotypically pluripotent (Fig. 2F, bottom half). However, as time progressed, their temporal trajectories diverged, coincident with differential cell fate acquisition (Fig. 2F, top half). Similarly, by plotting the trajectories of single-cells tracked early in differentiation as vectors on the embedding (Fig. 2G), we observed that individual cells’ trajectories started in the pluripotent area of the embedding and gradually diverged on the fate map as time progressed. Thus, multi-day, high-dimensional, image-based morphological profiling enhanced by deep learning allows to robustly map and predict cell fate evolution toward multiple, different cell fate outcomes, both at the population and at the single-cell level. We call our approach DEEP-MAP, for <u>deep</u> learning-enhanced <u>m</u>orphologic<u>a</u>l <u>p</u>rofiling.

### Proliferative features predictive of cell fate

With the DEEP-MAP NN model in place, we asked what features allows discrimination among different cell fate classes. Focusing on a small subset of biologically interpretable features, we carried out Bayesian network statistics and linear correlation analysis of the features and found that several features correlated directly with the cell fate class/label, indicating that they may be linked with the phenotypic states associated with the different cell fates (Fig. 3A). We then computed mutual information to quantitatively measure the mutual dependency between every feature and cell fate, as a way to estimate the importance of each feature to cell fate. Overall, we found that cell density and G1 status (as assessed by FUCCI-red signal intensity) were the features with the highest importance for predicting the different cell state classes (Fig. 3B), with different features displaying different degrees of importance across the different cell states. Specifically, cell density, G1 status and cell speed had high importance in both pluripotent and EC cells, with cell speed and nuclear area taking on increased importance in EC cells relative to pluripotent cells. These data suggest that cell speed and nuclear area are features that are discriminant of EC cells with respect to pluripotent cells, and that changes in cell migration control and shape/size may accompany pluripotency exit and onset of early ectodermal fate (Fig. 3C, top). By contrast, we found in ME cells that G1 status and cell speed had lower importance, and that the average distance of cells from the border of colonies (another measure of cell density) had a higher importance relative to pluripotent cells. This suggests that G1 status, cell speed and cell density discriminate ME cells, and that changes in cell cycle progression, migration and adhesion may accompany the onset of early mesodermal fate (Fig. 3C, bottom left). Finally, in EN cells we found that G1 status and cell speed had higher importance, and the average distance of cells from the border of colonies and cell density had a lower importance, relative to pluripotent cells, suggesting that those features are discriminant of EN cells and that changes in the control of cell cycle progression, migration and cell density are key at the onset of early endodermal fate (Fig. 3C, bottom right). Mapping of high importance features on the NN embedding (Fig. 3D) as well as overlaying them on image sequences (Fig. S3A-S3D and Movies S5-S6) confirmed visually that those features change coincident with fate changes. Hence, DEEP-MAP-derived phenotypic profiling information can be used to quantitatively identify, in an unbiased manner, morphological and proliferative properties predictive of cell fate transitions and point to possible mechanisms that cause or accompany commitment to different cell fates.

**Figure 3.**
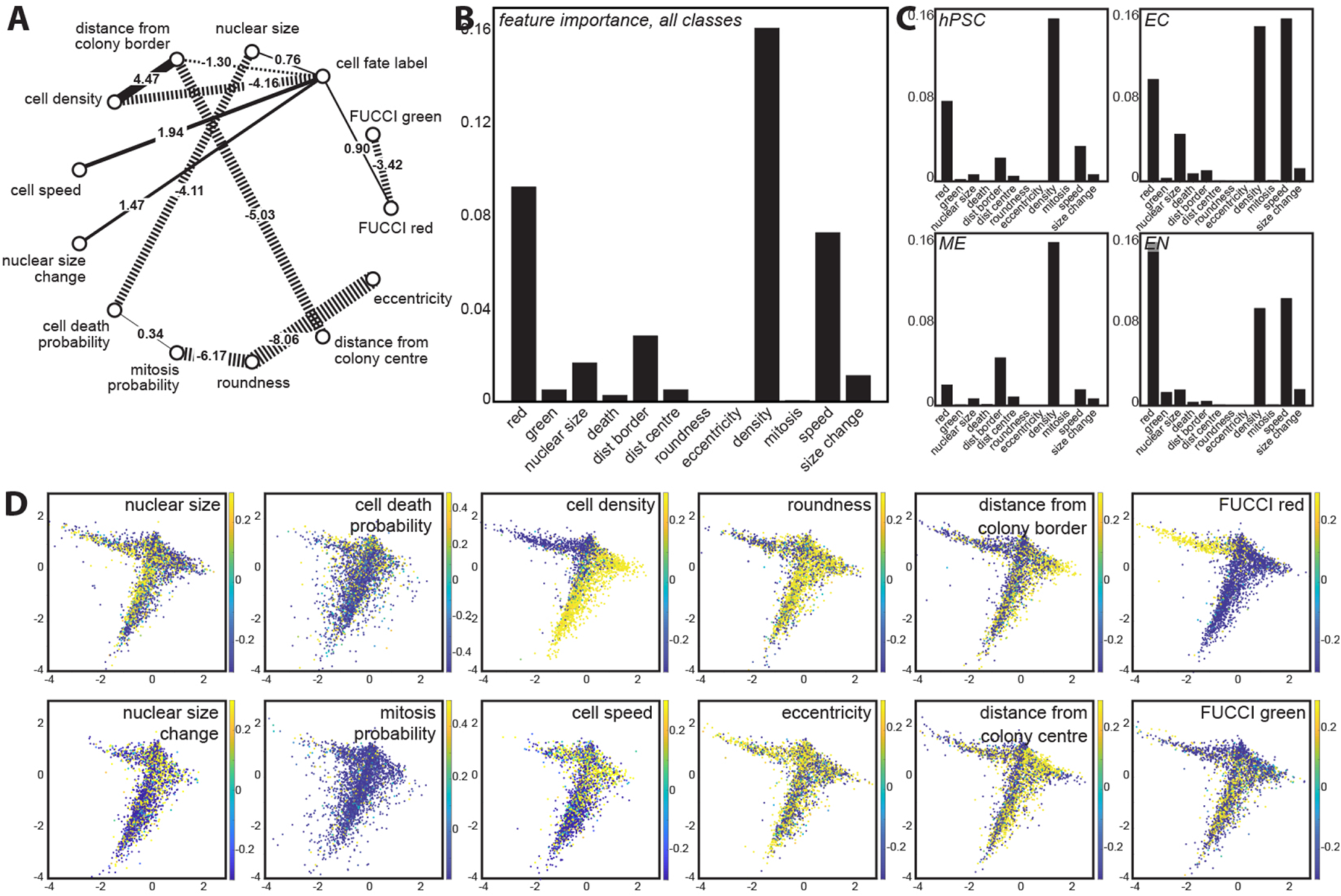
Identifying proliferative features predictive of different cell fates. **A**, Correlation coefficients and linkages between pairs of biologically interpretable features (solid line: positive correlation, dotted line: negative correlation) and with respect to cell fate label. As can be seen in the diagram, most features have direct correlations with the cell fate class/label. **B** and **C**, Mutual information analysis between biologically interpretable features and cell fate as a way to measure the importance of each feature in capturing the phenotypic state, for all classes together (**B**) versus each of the four cell states analysed separately (hPSC, EC, ME and EN; **C**). Cell density and G1 status (as assessed by FUCCI-red signal intensity) are the features with the highest importance for predicting the different cell state classes overall (**B**), with different features displaying high importance for different cell states. **D**, Mapping of biologically interpretable feature values on the neural network embedding, confirming that features predicted in **C** as being important for specific fate changes can be seen to also change accordingly on the embedding, demonstrating that dynamical morphological profiling can be used to quantitatively identify morphological and proliferative properties predictive of the different cell fate transitions.

### Proliferative state predicts single-cell fate dynamics, transition timing and fate intermediates

Next, we sought to exploit the predictive capacity of DEEP-MAP to observe single-cell fate dynamics in real-time. To this end, we used the NN cell fate class predictions to visually augment image sequences by displaying on the images the predominant (highest probability) predicted class for each single-cell through time. As expected, we observed that the overwhelming majority of pluripotent, EC, ME and EN triggered cells were predicted initially to be pluripotent (Fig. 4A-4D, left most image panels and Movies S7-S10), and that as time proceeded pluripotent cells maintained that predicted fate even as colonies grew larger and denser (Fig. 4A). By contrast, after the trigger, EC cells began losing the pluripotent phenotype ∼1 day later and began acquiring ectoderm phenotype by 2.5 days (Fig. 4B). Importantly, both of these timepoints are earlier than those at which pluripotency exit (e.g. OCT4 and NANOG loss, ∼2-3 days (Li et al., 2011)) and ectodermal fate onset (e.g. Nestin and PAX6 gain (Li et al., 2011) and SOX1 gain (Kim et al., 2021), ∼3-5 days) have been detected by quantitatively monitoring transcription factor levels. Hence, by using solely indirect proliferative readouts, DEEP-MAP is capable of detecting cell fate changes in ‘live’ cells earlier than what is currently technically possible. We were surprised to see that within only a few hours of receiving the differentiation trigger, ME and EN cells began losing the pluripotent phenotype (Fig. 4C-4D). ME cells began to acquire the mesoderm phenotype within 1 day of receiving the differentiation trigger and increasingly acquired that predicted fate for multiple days (Fig. 4C). By contrast, EN cells acquired the endoderm phenotype 2.5 days after trigger (Fig. 4D). Both of these differentiation onset timings are as early and possibly earlier than those at which early mesoderm (e.g. T gain, ∼1-2 days; (Rao et al., 2016)) and early endoderm (e.g. FOXA2 and SOX17 gain, ∼3 days; (Ogawa et al., 2013; Teo et al., 2011)) fate onset have been reported to occur. Taken together, these data indicate that DEEP-MAP can detect ‘live’ and dynamical cell fate-associated phenotypic changes earlier than previously possible, with unprecedented sensitivity and predictive power.

**Figure 4.**
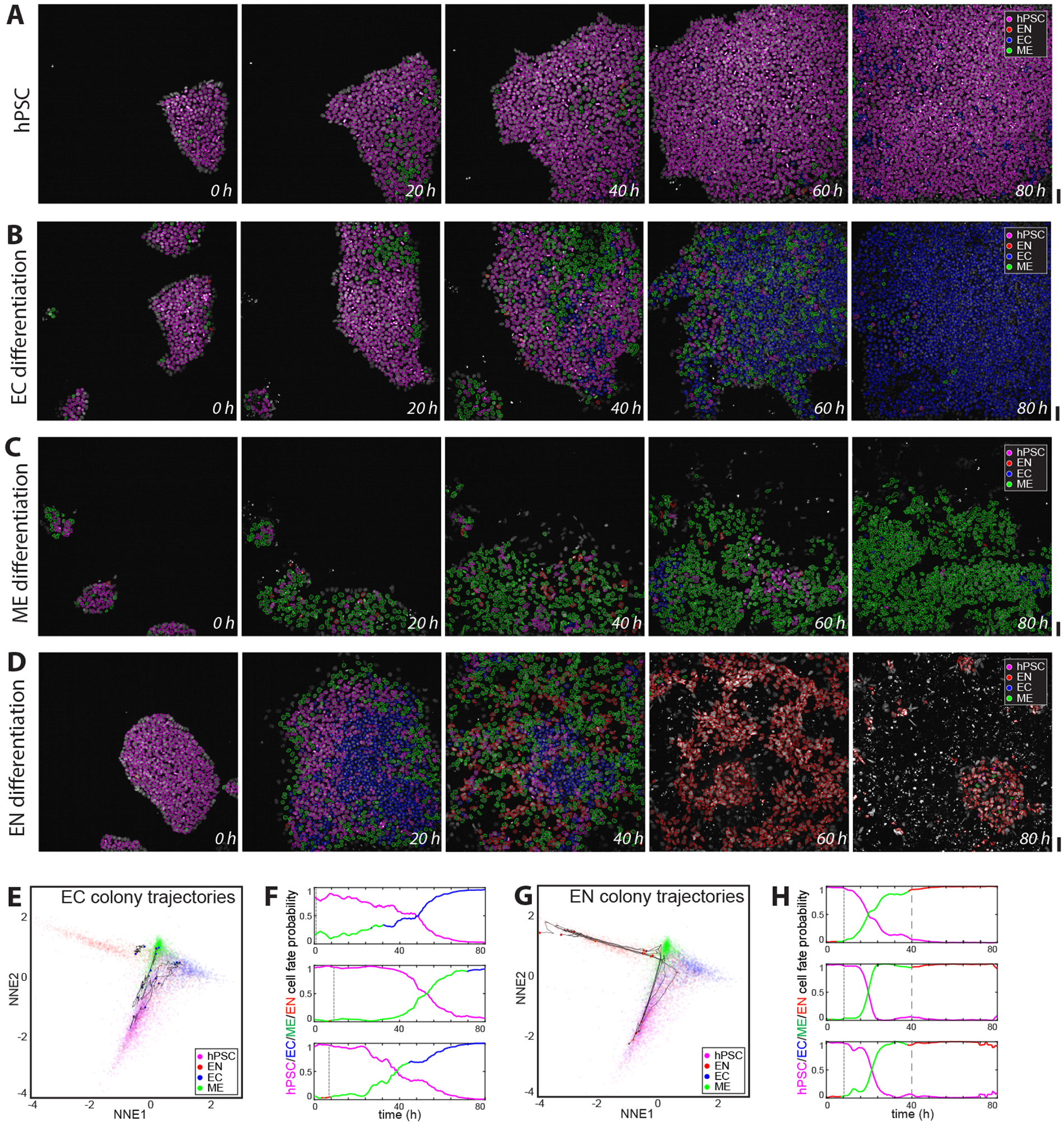
Morphological and proliferative state predicts cell fate dynamics and transitions in real-time. Image galleries of FUCCI and H2B-miRFP670 co-expressing cells imaged continually by optimised, multi-day time-lapse microscopy for 5 days as they either maintain pluripotency (**A**) or after receiving trigger for ectoderm (**B**), mesoderm (**C**) or endoderm (**D**) differentiation at day 0. FUCCI signal is not shown, only H2B signal is shown faintly in the images. Images are visually augmented by showing overlaid on the original images the predominant (highest probability) fate predicted by the DEEP-MAP neural network through time at single-cell level for each cell detected. Images are shown from the first 80 hours of time-lapse. Magenta/blue/green/red: predicted hPSC/EC/ME/EN fate, correspondingly (colour code shown as image inset). hPSCs are robustly predicted correctly as hPSCs through time regardless of changes in colony size and density (**A**), while EC, ME and EN gradually lose predicted hPSC status after just ∼1 day of differentiation trigger, and evolve differently toward different fates (**B**, **C** and **D**). **E**, Real-time phenotypic trajectories of different EC colonies (solid black lines corresponds to different colonies) showing most colonies’ trajectories beginning in the pluripotency domain (bottom centre of the plot) and evolving toward the EC domain (middle right of the plot) through time, with colonies showing visible heterogeneity in their trajectories. **F**, Plots showing the temporal evolution of the predicted cell fate assignment for three different EC colonies as a function of time. Magenta curves show the predicted probability of a colony having hPSC fate, as a function of time; blue/green/red curves show the predicted probability of the colony having EC/ME/EN fate respectively, with the colour displayed corresponding to the predominant (highest probability) fate only. As can be seen from the plots, EC colonies differ in their time of departure from pluripotency (dotted lines) as well as in the time of acquisition of the target EC (blue) phenotype, indicative of heterogeneity in the way the colonies acquire the target fate. **G**, Real-time phenotypic trajectories of different EN colonies showing most colonies’ trajectories beginning in the pluripotency domain and evolving toward the EN domain (middle right of the plot) through time, with colonies showing low heterogeneity in their trajectories. **H**, Plots showing the temporal evolution of the predicted cell fate assignment for three different EN colonies as a function of time. Colour codes as before. As can be seen from the plots, EN colonies departed from pluripotency with similar timing (dotted lines) and also acquired the target EN (red) phenotype with similar timing (dashed lines), suggesting a possibly tighter controlled response of cells and colonies to the EN differentiation triggers. In **G** and **H**, EN colony trajectories appear to take an intermediate ME phenotype (green) before acquiring their final EN phenotype, suggesting that an early ME-like state might be a fate intermediate during EN differentiation. Scalebars: 50 μm.

To gain further insights into the dynamics of lineage differentiation, we used the DEEP-MAP NN embedding to map real-time phenotypic trajectories of different EC colonies, by computing the average coordinates of each cell colony at every point in time through the 3.5 days of imaging (Fig. 4E). While most EC colonies’ trajectories began in the pluripotency domain and later evolved toward the ectoderm domain, they appeared to do so heterogeneously. By computing the predominant (highest probability) cell fate assignment for each separate colony as a function of time, we confirmed that colonies differed in the time of departure from pluripotency as well as in the time of acquisition of the majority ectodermal phenotype (Fig. 4F). By contrast, we found that EN colonies’ trajectories proceeded much more homogeneously through the DEEP-MAP fate map, suggestive of a much more tightly controlled response of cells and colonies to differentiation triggers (Fig. 4G). Strikingly, all EN trajectories appeared to go through the mesoderm domain en route to acquisition of the endoderm phenotype, suggesting that acquisition of an early mesoderm-like state could be an intermediate in acquiring endodermal fate. This was evident when we looked at the colonies’ predominant cell fate assignment as a function of time, which showed that EN-triggered colonies departed pluripotency early and almost synchronously, then acquired and maintained a predicted mesodermal phenotype for 1-1.5 days, and subsequently acquired an endodermal phenotype 40 hours later (Fig. 4H). Altogether, our findings demonstrate that, by integrating high-dimensional, image-based morphological profiling with deep learning, DEEP-MAP can predict cell fate dynamics ‘live’ in real-time with high temporal sensitivity across multiple fates. Hence, this technology can be leveraged to reveal differences in cell fate dynamics between lineages, cells and colonies and can reveal the history, timing, and existence of cell fate intermediates, which can be elusive to fixed-cell technologies.

## DISCUSSION

We have established DEEP-MAP, a deep learning-enhanced morphological profiling approach, which enables ‘live’ large-scale microscopy imaging and phenotyping of hPSC populations at single-cell level and predicts cell fate dynamics and transitions in real-time over several days. We generated by CRISPR knock-in hPSC lines stably co-expressing ‘live’ cell proliferation reporters and then established multicolour time-lapse imaging compatible with long-term cell viability and health, and compatible with proliferation and differentiation under the microscope. Our pipeline integrates image processing, machine learning and statistical analysis workflows to enable time-resolved, high-dimensional phenotyping of 100’000s of single-cells, as well as deep learning methods allowing robust visualisation, mapping, phenotypic clustering and prediction of cell fate dynamics toward multiple fate outcomes. We show that based on two very general and commonly used cell proliferation reporters - FUCCI and fluorescent H2B – DEEP-MAP yields rich, deep and continuous information about morphological and proliferative state that can predict and reveal cell fate dynamics ‘live’ in real-time and at single-cell level without a need for customised cell state reporters specific to the fates and lineages being monitored. The fact that the reporters used here are universally present and visible across developmental and cell differentiation contexts makes our approach highly versatile and broadly applicable.

Our choice to use nuclear-localised fluorescent reporters for morphological profiling was not fortuitous: hPSC colonies are very compact and therefore we used nuclear reporters to be able to unequivocally detect and, where possible, track cells through time. However, DEEP-MAP could be adapted to use other ‘live’ Cell Painting-type (Bray et al., 2016) morphological reporters, both genetically encoded and membrane permeable (e.g. fluorogenic probes (Mishra et al., 2019)), and applied to other cell types of choice. Similarly, DEEP-MAP could form the basis of methodologies to carry out label-free cell fate mapping and prediction (Christiansen et al., 2018; Ounkomol et al., 2018), by applying it to cells that are spatially sparse and can be unequivocally identified without the need for fluorescent reporters (Al-Zaben et al., 2019) or by using label-free imaging optical modalities (Kallepitis et al., 2017; Marrison et al., 2013).

Previous efforts have used deep learning to detect and predict cell states ‘live’ before. For instance, convolutional neural networks (CNNs) can detect morphological differences between unlabelled pluripotent and early epiblast-like differentiating cell patches of mouse embryonic stem cells just 20 minutes after onset of differentiation (Waisman et al., 2019). Similarly, CNNs and recurrent neural networks (RNNs) can predict lineage choices of individual, disaggregated primary mouse hematopoietic stem and progenitor cells (HSPCs) up to three generations before molecular marker annotation, based on their morphological and displacement characteristics (Buggenthin et al., 2017). By combining morphological profiling applied to simple nuclear-localised cell proliferation readouts with supervised neural networks, DEEP-MAP goes significantly further. Our method captures the evolving dynamics of cell fate transitions over several days across entire colonies of highly compact human stem cells, demonstrating the suitability of our approach to large-scale, tissue-level fate dynamics’ prediction with single-cell resolution.

Using DEEP-MAP we found that, upon receiving a differentiation trigger, hPSCs initiate pluripotency exit between 8 and 24 hours. This is as early and possibly earlier than previously detected using ‘live’ fluorescently-labelled transcription factor reporters in hPSCs (Kim et al., 2021; Wolff et al., 2018). Furthermore we found that early endoderm-triggered colony trajectories appear to transiently go through a predicted mesoderm phenotype, suggesting that acquisition of an early mesoderm-like fate could be an intermediate in acquiring endodermal fate. This is in agreement with previous observations in mouse embryonic stem cells (Kubo et al., 2004; Mahmood and Aldahmash, 2015; Tada et al., 2005), and suggests that our approach can reveal the existence of cell fate intermediates that would be invisible without real-time phenomics information.

In addition, our analysis also resolved visible differences in the real-time cell fate dynamics within and between early EC and EN lineages. This suggests that DEEP-MAP could become a general framework to quantify the heterogeneity, noise and speed of different cell fate transitions. Accordingly, DEEP-MAP could help identify the origins of variability in synthetic tissue generation, as well as give insights into how to engineer tissues predictably and robustly (Brassard and Lutolf, 2019; Prochazka et al., 2017).

In sum, DEEP-MAP provides a framework to quantitatively investigate the dynamics of cell fate decisions at single-cell level within tissues in real-time and at scale. By enabling the generation of spatiotemporal, predictive maps of tissue formation, this approach can help to measure, benchmark and, ultimately, predict and control complex tissue design.

### Limitations of Study

In this study we used differentiation triggers known to result in high cell fate conversion rates. This allowed us to assume at the endpoint that all cells acquired pseudo-homogeneously the intended fates. This assumption simplified the deep learning strategy, as it enabled us to use a common neural network architecture to both learn to spatially map cells as well predict cell fate dynamics. Such a strategy - where we constrain the network to simultaneously learn cell fate prediction while generating a spatial map - could be limiting when investigating more complex tissues or heterogenous cell populations, where differentiation may be less efficient or specific and might lead to generation of a variety of differentiation products (including cells of unknown fate). In that case using separate network architectures to learn mapping and visualisation could significantly help increase the cell fate predictive power required by the biology. In addition, combination of the approach described here with spatially resolved transcriptomics approaches (Marx, 2021) might also become possible in the future, providing a way to obtain a more detailed landscape of the resulting cell types generated, as well as quantitative, deep readouts on the transcriptional status of cells. Such a combined approach could lead to the generation of ‘time machine’-type models of cell fate dynamics with increasingly precise, diverse, temporally-resolved and sensitive predictive power.

## STAR METHODS

### Construction of plasmids for CRISPR/Cas9 mediated knock-in

Each pair of sgRNAs to target Rosa26 locus cloned into All-In-One (AIO) CRISPR/Cas9 nickase plasmids (#74119, Addgene). Sequence of sgRNA 1 and sgRNA 2 is 5’- GTCGAGTCGCTTCTCGATTA-3’ and 5’-GGCGATGACGAGATCACGCG-3, respectively. For donor constructs of H2B-miRFP670 and FUCCI, the fragments of 5’ and 3’ homology arms of ROSA26 locus were subcloned into H2B-670 (modified from pmiRFP670-N1, #79987, Addgene) and FUCCI (kind gift from Ludovic Vallier’s lab, U. of Cambridge). All of the cloning procedures were performed using In-fusion HD Cloning kit (639650, Takara) for seamless DNA cloning.

### Generation of FUCCI/H2B-miRFP670 reporter hESCs lines

To establish hESCs that contain FUCCI (Sakaue-Sawano et al., 2008) and H2B (Kim et al., 2021; Shcherbakova et al., 2016) reporters, reporter constructs were introduced to one of the genomic safe harbour regions, *ROSA26* locus by using CRISPR/Cas9 nickase to minimise insertional mutagenesis. Cells were transfected with both AIO CRISPR/Cas9 nickase and donor vectors with ROSA26 homology arms using lipofectamine stem transfection reagent (STEM00008, Thermo Fisher Scientific) according to the manufacturer’s protocol. Briefly, 1 μg of each plasmid, total 2 μg was diluted into 7.5 μl of reagent in the 200 μl of Opti-MEM I medium (31985062, Thermo Fisher Scientific) and incubated for 10 min at RT. Transfected cells were sorted on a BD Influx and collected into 1.5 ml microcentrifuge tubes containing 500 ul of Knock-out serum replacement (10828028, Thermo Fisher Scientific). hESCs were maintained under essential 8 (A1517001, Thermo Fisher Scientific) medium on Geltrex (A1413301, Thermo Fisher Scientific)-coated plates and changed the medium every day.

### Differentiation of hESCs into multiple lineages

To differentiate into neuroecto lineages, hESCs were applied to PSC neural induction medium (A1647801, Thermo Fisher Scientific) according to the manufacturer’s instruction. Briefly, cells were grown in the induction medium (Neurobasal medium containing 50x PSC supplement) for 5 days of time-lapse experiment and the medium changed every other day. To differentiate into mesodermal lineage, hESCs were applied to PSC cardiomyocyte differentiation kit (A2921201, Thermo Fisher Scientific) according to the manufacturer’s instruction. Briefly, cells were grown in cardiomyocyte differentiation medium A for the first 2 days then it was changed into cardiomyocyte differentiation medium B for 3 days. Medium changed every other day. To differentiate into definitive endodermal lineages, cells were applied PSC definitive endodermal induction kit (A3062601, Thermo Fisher Scientific) according to the manufacturer’s instruction. Briefly, cells were grown in PSC definitive endoderm induction medium A for 1 day, then changed to definitive endoderm induction medium B till the end of experiment.

### Time-lapse imaging

Established FUCCI/H2B hESC lines were plated onto Geltrex-coated CellCarrier-96 Ultra Microplates (6055302, Perkin Elmer) a day before imaging. Differentiation trigger was applied by gently changing to the differentiation medium before imaging. Cells were imaged using a Yokogawa CV7000 high throughput confocal microscope (Wako). FUCCI signal was captured every 30 min using both 488 and 561 nm at 150 and 350 ms of exposure time and H2B-miRFP670 signal was captured using 640 nm at 500 ms of exposure time every 5 min, respectively. Time-lapse imaging was performed for 5 days.

### LEVER Processing

The open source LEVER software package (Cohen, 2014; Wait et al., 2014; Winter et al., 2016) (https://leverjs.net) is utilised to segment and track cells from hPSC time-lapse image sequences. The H2B channel TIFF files generated from the microscopy experiments are imported into the LEVER file format, and an ensemble-based segmentation algorithm and a cell tracking algorithm are then applied to the image sequences. The segmentation algorithm separates foreground and background regions through combining an adaptive intensity thresholding with a Laplacian of Gaussian filter. The only parameter to the algorithm is the minimum cell radius, here set at 2.5μm. Following the segmentation, cells are tracked and mitotic events identified as described previously (Winter et al., 2015).

### Feature Extraction

Feature extraction is carried out using a custom script MATLAB (MathWorks). The segmentation results generated by LEVER are read into MATLAB and correlated with the original TIFF images generated by the microscopy experiment across the H2B and FUCCI channels. The script then advances frame by frame, using the segmentation results as a registration method to record features from the original images. Feature extraction only occurs at timepoints where all 3 channels are present. Images undergo local background subtraction using a 50x50 pixel sized rolling average filter in order to correct for spatial variations in microscopy illumination.

Colonies of cells are identified using the DBScan algorithm (Sander et al., 1998) and tracked through time using shared cell identities between time frames. The script iterates through each colony and then through each cell belonging to that colony in order to extract a suite of numerical features. Features extracted include: fluorescence distribution features, texture features such as the Haralick Features (Haralick et al., 1973), Hu’s Invariant Moments (Flusser, 2000; Marchant, 2021; Ming-Kuei, 1962), Zernike moments (Saki et al., 2013; Tahmasbi et al., 2011), Gray Level Run Length Matrices (Galloway, 1975; Wei, 2021), Gray Level Size Zone Matrices (Thibault et al., 2013), Neighborhood Grey Tone Difference Matrices (Amadasun and King, 1989; Vallières et al., 2015), shape descriptors, colony based features such as distance from the colony centroid and border, and cell density features. Haralick features are calculated by cropping each segmented cell and calculating the texture features of the cropped image. The Haralick features calculated for the H2B channel are used to predict the cell (i.e. nuclear or chromatin) state of a cropped cell using a trained Support Vector Machine (Allwein et al., 2000). The Support Vector Machine assigns the most likely class to each segmented cell from 5 different classes: ‘interphase’, chromatin is decondensed and occupies a mostly round nucleus, implying that the cell is in G1, S or G2 phases of the cell cycle; ‘metaphase’, condensed chromatin with chromosomes aligned on a metaphase plaate; ‘anaphase’, two nearby masses of condensed chromatin corresponding to the cell having undergone the metaphase to anaphase transition; ‘apoptosis’, nuclear debris, i.e. fragments of apparently condensed chromatin typical of cell death; and ‘mis-segmented’, the H2B signal segmentation is poor and corresponds possibly to more than one cell’s nucleus (Note: this latter class is not shown in Figure 1 or mentioned in Figure 1 legend for simplicity). The SVM was trained manually with 500 cells in each class and has a prediction accuracy on the training set of >90% across all classes. Dynamical features such as cell speed, and cell size change are calculated by referencing the previous frame where a cell is detected and calculating the difference in features between the two timepoints.

Extracted features are stored in an SQLite database for downstream processing.

### Feature Pre-processing

Dynamical features are not present in a significant proportion of data points as cell tracking does not occur at 100% efficiency. As a result, the stored database contains NaN values for certain features, which the downstream Neural Network deals with poorly. To prevent failure in the Neural Network, features that commonly contain NaN values such as cell speed and cell size change are filtered out, as well as any rare individual data points that have NaN values due to failures in calculating other features. Other features such as cell XY coordinates, the colony identity, and parameters used to calculate shape features that are deemed unnecessary for cell fate prediction are also removed. This filtering step filters the number of available features for the neural network (NN; see later for NN implementation) to train on, from the 603 originally recorded features to 564 features used for actual training and embedding. The filtered data points are assigned labels according to which experimental condition the datapoint is derived from, in order to facilitate downstream training of the NN.

### Neural Network Embeddings

After feature pre-processing, datasets are fed into the NN embedding pipeline (see later), generating an XY coordinate for each datapoint that is recorded along with the cell ID, frame, label, and classification probabilities of belonging to each differentiation class. Neural network embeddings are plotted using MATLAB using the scatter function, with the data points being coloured according to the recorded dataset label. Real time cell population trajectories are plotted by calculating the mean XY coordinates at each timepoint for a given experimental condition. The mean XY coordinates are then averaged using a rolling window of 2.5 hours to reduce the effect of random noise and fluctuations in the dataset. The averaged population trajectory is plotted over the embedded datapoints with markers indicating the starting and ending time point. Vectorial single cell trajectories are calculated by identifying the 20 longest sequential tracks of cells generated by the LEVER tracking algorithm for each experimental condition. For these 20 tracks, an arrow is plotted from the start to the endpoint of that cell’s tracked trajectory. Proliferative feature distributions are displayed on the embedding plot by changing the method of colouring individual embedded data points to a chosen extracted feature. Features chosen to be plotted are pulled from the extracted feature database and correlated with the XY embedding coordinates using the stored cell ID and frame information. The features are standardised using the Z-score method and the selected feature is used to individually colour the scatter points.

### Visualising changes in proliferative features and cell fate

Pseudocolor images displaying the proliferative features for each cell are generated using the stored segmented cell boundaries generated by the LEVER segmentation package. The cell segments generated by LEVER are correlated with the extracted feature data points for an individual time point. The cell segments are displayed in a MATLAB figure, coloured using a selected proliferative feature, and then saved to generate sequential images of proliferative features changing through time. Changes in cell fate are visualised using the classification probabilities generated by the NN embedding, with the original microscopy images of the H2B channel and the segments generated by LEVER. The microscopy image is displayed in MATLAB, and the segments generated by LEVER are plotted and overlayed over the microscopy image. The colour of the segmentation outline is determined by the cell fate class with the highest classification probability as calculated by the NN classifier. The resulting image is written into a TIFF file and sequential frames are concatenated to the TIFF file to generate an image stack that displays the cell fate transitions of the experimental population through time.

### Cell Fate Probability Tracks

Cell fate probability tracks for individual colonies are plotted using the classification probabilities assigned to each data point in that colony by the NN embedding. To compare the level of differentiation each colony has undergone, the classification probabilities of the non-pluripotent states are summed together to represent the degree of differentiation. The classification probabilities of all data points at a given frame within a colony is averaged together and the mean probability is plotted over time. The differentiated probability plot is coloured according to which differentiated state has the highest classification accuracy at that given frame. A rolling average window of 2.5 hours is applied to the classification probabilities in order to minimise the amount of noise in the trajectories.

### Embedding and prediction by Deep Neural Network

#### Overview of the idea

Given the poorly separable embedding obtained by the unsupervised machine learning (ML) algorithms namely PCA (Abdi and Williams, 2010), t-SNE (van der Maaten and Hinton, 2008) and UMAP (McInnes et al., 2018), we devised a supervised deep learning approach capable of extracting an embedding and performing predictions. The advantage is two-fold, the DL model’s powerful representation boosted by the guiding label information brought by the supervised learning approach.

Although our framework is pipeline-based involving multiple stages with engineering-based feature extraction, the DL embedding model refines the extracted features by learning informative/discriminative representations. Its multi-layers progressively reduce the features’ dimensionality with each layer as it learns higher-levels of feature abstraction and finally outputs a two dimensional representation at the last layer. This final representation serves as our 2D visualisable embedding of the input features.

Unlike unsupervised DL embedding extraction methods (David and James, 1987; Vincent et al., 2008) that use reconstruction loss for training, our loss exploits the available label providing sharper training guidance. That is, our learning approach is fully supervised with labels used to project the embedding of different classes in different separable clusters/regions. This is done by replacing the standard hyper-plane classification layer by a clustering classification one. That is, a cluster is assigned to each class and the input’s class is predicted with probability proportional to its distance from the clusters’ centre. This alters the loss function to penalise clusters close to each other and embeddings not falling in its assigned cluster (more details further below).

#### Detailed Pipeline

Fig. 2D shows the DL-based embedding pipeline, which can be divided into three stages. First, the raw input image data are segmented and tracked using LEVER (see LEVER Processing section) to produce cell patches, which are further processed to extract morphological, fluorescence intensity and texture features at single-cell resolution (see Feature Extraction section). We then established a multi-layer DL NN to learn an informative/discriminative representation from the features. Although DL can be used to learn features from raw data in a fully automated way without experts’ involvement, in our case, we used DL to refine the features by further reducing the dimensionality and noise before generating a 2D embedding at its last layer.

As stated above, training this embedding pipeline was done in a supervised way where labels were used to establish the objective function of the optimisation problem. To enforce separable embedding outputs in cloud shape, we replaced the classification layer normally attached to the NN by a clustering one. Nevertheless, our embedding-clustering network can still provide predictions as well as an embedding representation. In the following, we provide details of our NN architecture including both the embedding and the clustering components.

#### Network architecture

Our embedding network consists of 5 fully connected layers (the number of layers was chosen based on optimal performance), each followed by a ReLu non-linear function. The first layer reduces the input dimensions to hidden_dim dimensions that are then progressively reduced by 2 and finally to 2D at the last layer. Thus, the outputs of layers 2,3,4, and 5 are hidden_dim/2, hidden_dim/4, hidden_dim /8 and 2 respectively. These multi-layers of non-linear transformations represent multi-levels of automatic feature extractions, where the input features at each layer are further refined so as to end up with an optimal features’ output for the purpose of embedding representation expressed in the objective function. Denote ***e*** ∈ *R*^2^ the embedding of input *x* ∈ *R*^n^ where *n* is the number of input features. We express the embedding network transformation by *ϕ*_*w*_ governed by the network layers weight *W*. The embedding ***e*** = *ϕ*_*w*_ (*x*) are then fed to the clustering network serving two purposes, prediction and to build the learning objective function. To force the embedding of a given class to fall in an isolated cloud, the clustering layer assigns a cluster centre to each class and expresses a probability distribution for each embedding over the classes, which is proportional to the distance from the classes’ centres. Denote *o*_*i*_ ∈ *R*^2^ the centre of the cluster corresponding to class *i*, thus, the prediction *y* ∈ {1, …, *C*} of embedding ***e*** is distributed according to a categorical distribution governed by parameters *p* = (*p*_1_, … *p*_c_) where *C* is the number of classes and *p*_1_ ∝ 1 - ||*c*_1_ - *e* || such that 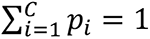. To meet this probability condition (*p*_i_≥ 0 ∀*i* ∈ {1, …, *C*} 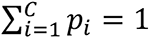, we utilise a Softmax non-linear function on the distance vector from the embedding to all centres (||*o*_1_ − ***e***||, . . . ||*o*_c_ − ***e***||), 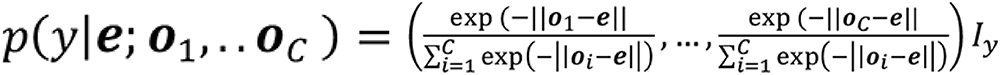 where *I*_y_ ∈ {0,1}^c^ is a vector with all dimensions set to zeros except that at position *y* is set to one. Although norm 2 function penalisation of the embedding quadratically diminishes as it gets closer to its corresponding centre, an even more relaxed penalisation around the centre similar to that of SVM hyperplane soft separation would result in more aesthetically satisfying embedding. This relaxation can be achieved by adding a nonlinear function -tanhshrink (≡ - *tanhs(x) + tanhs(x) - (x)* before the softmax function, thus the parameters of the categorical probability distribution over classes become, 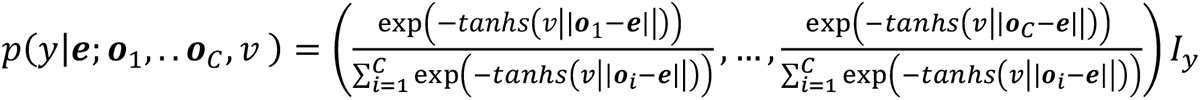 where parameters *ν* controls the relaxation margin. Given the probability distribution of a given embedding over the classes, we can now either sample the prediction 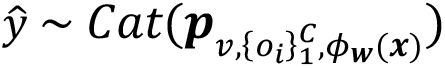 or take an argmax over the classes probabilities 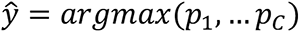, where 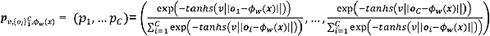.

#### Training method

So far, we have focused on the inference/feed-forward part of the model assuming the embedding parameters *w* and the clustering ones *o*, ν fixed. Next, we present the learning part (third stage in Fig 2D) of the method. As stated above, our approach is supervised, that is the ground truth (labels) are presented during learning. Thus, considering the model parameters as variables, we can use cross entropy as a loss function between the ground truth and the prediction outputs. Cross entropy loss is useful for training classification problems when the predicted output is governed by a multinomial/categorical distribution: 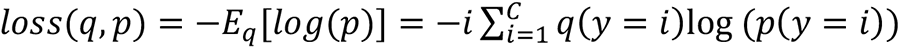 where #x1D45E; is the true probability. Since, our ground truth is deterministic, we can write the loss as − 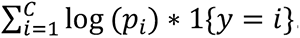. Having defined the objective function, the learning is achieved by solving this optimisation problem with respect to embedding and clustering parameters. We used the Adam optimiser (Kingma and Ba, 2014) with weight decay regularisation.

#### Datasets and Implementation

We split our dataset into three portions, 80 % for training, 15 % for validation and 5 % for testing. The validation is used for parameter tuning. We performed a grid search over different learning rates, weight decays, number of network layers, hidden dimensions and batch size. The parameters that gave the best results on the validation set were taken, namely, 0.0001 learning rate, 0.0001 weight decays, 512 hidden dimension, 64 batch size and 5 total fully connected layers. Our embedding-prediction code is implemented in Python 3.8 and using Pytorch 1.6. We trained on RTX 2080TI GPU on 53,120 (80 % of 66,400 total datapoints) datapoints consisting of four classes 20,933 Pluri, 18,927 Ecto, 1,036 Endo and 2,222 Meso. The quantitative prediction results on the testing set as well as on data from an independent biological experiment are shown below:

| Datasets | precision | recall | f1-score | # of points | Accuracy |
| --- | --- | --- | --- | --- | --- |
| Testing set from training experiment. | 0.85 | 0.89 | 0.87 | 10403 | 0.85 |
| Pluri | 0.88 | 0.71 | 0.79 | 1749 |  |
| Ecto | 0.85 | 0.83 | 0.84 | 9382 |  |
| Endo | 0.86 | 0.87 | 0.86 | 8465 |  |
| Meso |  |  |  |  |  |
| Dataset form new experiment |  |  |  |  | 0.68 |
| Pluri | 0.72 | 0.90 | 0.80 | 206945 |  |
| Ecto | 0.85 | 0.31 | 0.46 | 158067 |  |
| Endo | 0.69 | 0.75 | 0.71 | 198148 |  |
| Meso | 0.48 | 0.67 | 0.56 | 80491 |  |

### Correlation and features importance by Bayesian statistics

Given the black-box nature of NN outputs, we sought to recall biological interpretability for our prediction/embedding pipeline. Supported by the pipeline’s hybrid approach involving both manual feature extraction and DL feature learning components, we employed Bayesian statistics to provide declarative representation of how the input extracted features interact with each other. For this analysis, we only selected biologically interpretable features.

The methodology is divided into two main steps. Firstly, we learn a Bayesian network (i.e, Directed Acyclic Graph (DAG) representation) (Koller and Friedman, 2009) consisting of nodes representing the biologically interpretable features as random variables and edges expressing the conditional independency among these features. Doing so makes it computationally feasible to perform/approximate inference across the variables. Secondly, we use the trained network to perform inference, allowing us to compute linear correlations (Benesty et al., 2009) among the variables as well as the mutual information with the different fates.

The step of learning the Bayesian network is divided into two tasks. First, we learn the Bayesian network structure from the data using Hill-Climb Search (Koller and Friedman, 2009). The learned network structure expresses the conditional independency among features, but it does not encompass the optimal numerical parameters needed to compute the joint distributions and perform inference. To learn the network parameters, we maximize the expected log likelihood (Koller and Friedman, 2009) of the joint distribution defined by the learned network structure over the data with respect to its parameters.

Having learned the network structure and parameters, we are ready to infer needed information about the features’ correlations as well as their importance to the cell fate. Fig 2A shows linear correlation among all interpretable features plus the cell fate. To compute this correlation, we first convert the Bayesian network to a Markov network using moralization (Koller and Friedman, 2009), and we then compute the covariance matrix using the marginal distribution induced by the network and take the values of connected variables. Linear correlation shows how features interact; however, they do not express the features importance to the cell fate. We used mutual information as a metric to measure the mutual dependency between every feature and the cell fate. The lower the mutual importance, the more independent the feature is from the cell fate, implying that less information is conveyed by the feature to the cell fate. That is, it expresses the feature’s importance to the cell fate. Computing the mutual information between two variables entails knowing the joint and marginal distribution of these variables. We used Belief Propagation method to compute these distributions. Fig. 2B presents this feature importance measure for all the biologically interpretable features. In this figure, we do not set the cell fate variable and measure the importance to cell fate in general. In Fig. 2C, we set the cell fate variable as evidence and provide the feature importance to each of the four cell fate classes. The Bayesian analysis code is implemented in Python.3.8 using the pgmpy 0.1 package. We adopt discrete Bayesian Network whose structure and parameters are learned using 391,725 discretised datapoints. Discretisation was performed using sklearn pre-processing package.

## ACKNOWLEDGEMENTS

We thank John Gurdon, Tony Kouzarides, Ludovic Vallier, Daniel Ortmann, Jose Silva, Laura Wagstaff, Paola Marco Casanova, Anatole Chessel, Balint Antal and Faraz Janan for early experimental/computational support or discussions, Lucia Marucci, Martin Homer, Martin Hemberg and members of the Carazo Salas group for feedback, C. Lilliehook (Life Science Editors) for editorial services, and Andrew Herman, Lorena Sueiro Ballesteros and the University of Bristol Flow Cytometry Facility for help with transgenic cell line generation. This work was supported by an EPSRC PhD studentship (E.R.), a Wellcome Trust PhD studentship (S.F.H.), a China Scholarship Council PhD studentship (Y.S.), Human Frontier Science Program (HFSP) grant RGP0043/2019 (R.E.C.S., A.R.C.), University of Bristol funds (R.E.C.S.), Cancer Research UK Programme Foundation Award C38607/A26831 (E.P.) and Wellcome Trust Senior Research Fellowship 205010/Z/16/Z (E.P.).

## AUTHOR CONTRIBUTIONS

R.E.C.S. conceived the research and designed and supervised the study with help from E.P and A.R.C. S.F.H, E.R. and S.K. together established optimised conditions for multi-colour, time-lapse high-content hPSC imaging compatible with multi-day, healthy cell proliferation and differentiation. S.K. carried out reporter and vector construction, reporter knock-in and transgenic multi-reporter cell line generation. S.K. and E.R. carried out multi-day time-lapse microscopy experiments. E.R. did quantitative image and data analysis in MATLAB and R, and image analysis and cell tracking using LEVERJS with continuous help from A.R.C. Y.S. contributed with image analysis quality control. S.M. established neural network embedding methods and predictive cell fate models, with feedback from E.R. and R.E.C.S. R.E.C.S. wrote the paper with help from E.P. All authors read and commented on the manuscript.

## SUPPLEMENTAL INFORMATION

### SUPPLEMENTAL VIDEO CAPTIONS

#### Supplemental Video 1

3.5 day time-lapse video sequence of a hPSC cell line co-expressing FUCCI and H2B- miRFP670, generated by CRISPR knock-in. FUCCI and H2B-miRFP670 co-expressing hPSCs are cultured in pluripotency maintaining conditions. To minimize phototoxicity to cells, an optimised imaging modality was used where H2B-miRFP670 signal was captured every 5 minutes – to enable continued cell/colony detection and tracking - and FUCCI signal every 30 minutes.

#### Supplemental Video 2

3.5 day time-lapse video sequence of FUCCI and H2B-miRFP670 co-expressing hPSCs, trigggered to undergo ectoderm (EC) differentiation at time 0. H2B-miRFP670 signal was captured every 5 minutes and FUCCI signal every 30 minutes.

#### Supplemental Video 3

3.5 day time-lapse video sequence of FUCCI and H2B-miRFP670 co-expressing hPSCs, trigggered to undergo mesoderm (ME) differentiation at time 0. H2B-miRFP670 signal was captured every 5 minutes and FUCCI signal every 30 minutes.

#### Supplemental Video 4

3.5 day time-lapse video sequence of FUCCI and H2B-miRFP670 co-expressing hPSCs, trigggered to undergo endoderm (EN) differentiation at time 0. H2B-miRFP670 signal was captured every 5 minutes and FUCCI signal every 30 minutes.

#### Supplemental Video 5

3.5 day time-lapse video sequence of hPSCs triggered to undergo ectoderm (EC) diffferentiation at time 0, corresponding to the same EC cells and colonies shown in Figure S1. Images are fake coloured to display the feature value levels for cell density - a feature shown in Figure 3 to have high importance in distinguishing different cell fates - through time at single-cell level for each cell detected. Note that EC cells show high density through time.

#### Supplemental Video 6

3.5 day time-lapse video sequence of hPSCs triggered to undergo endoderm (EN) diffferentiation at time 0, corresponding to the same EN cells and colonies shown in Figure S1. Images are fake coloured to display the feature value levels for cell density - a feature shown in Figure 3 to have high importance in distinguishing different cell fates - through time at single-cell level for each cell detected. Note that in EN cells cell density changes dramatically through time.

#### Supplemental Video 7

Visually augmented 3.5 day time-lapse video sequence of hPSCs cultured in pluripotency maintaining conditions, displaying predicted single-cell fate in real-time. The sequence corresponds to the the same pluripotent cells and colonies shown in Figure S1 and Video S1. The cells’ FUCCI signal is not shown, only H2B signal is shown faintly in the images, which instead show overlaid on the original image sequences the predominant (highest probability) fate predicted by the DEEP-MAP neural network through time at single-cell level for each cell detected. Magenta/blue/green/red: predicted hPSC/EC/ME/EN fate, correspondingly (colour code shown as image inset). hPSCs are robustly predicted correctly as hPSCs through time regardless of changes in colony size and density.

#### Supplemental Video 8

Visually augmented 3.5 day time-lapse video sequence of ectodermal-triggered hPSCs, displaying predicted single-cell fate in real-time. The sequence corresponds to the the same ectodermal (EC) triggered cells and colonies shown in Figure S1 and Video S2. The cells’ FUCCI signal is not shown, only H2B signal is shown faintly in the images, which instead show overlaid on the original image sequences the predominant (highest probability) fate predicted by the DEEP-MAP neural network through time at single-cell level for each cell detected. Magenta/blue/green/red: predicted hPSC/EC/ME/EN fate, correspondingly (colour code shown as image inset). EC triggered cells lose predicted hPSC status after just ∼1 day following differentiation trigger and evolve gradually toward the EC fate.

#### Supplemental Video 9

Visually augmented 3.5 day time-lapse video sequence of mesodermal-triggered hPSCs, displaying predicted single-cell fate in real-time. The sequence corresponds to the the same mesodermal (ME) triggered cells and colonies shown in Figure S1 and Video S3. The cells’ FUCCI signal is not shown, only H2B signal is shown faintly in the images, which instead show overlaid on the original image sequences the predominant (highest probability) fate predicted by the DEEP-MAP neural network through time at single-cell level for each cell detected. Magenta/blue/green/red: predicted hPSC/EC/ME/EN fate, correspondingly (colour code shown as image inset). ME triggered cells lose predicted hPSC status after just ∼1 day following differentiation trigger and evolve gradually toward the ME fate.

#### Supplemental Video 10

Visually augmented 3.5 day time-lapse video sequence of endodermal-triggered hPSCs, displaying predicted single-cell fate in real-time. The sequence corresponds to the the same endodermal (EN) triggered cells and colonies shown in Figure S1 and Video S4. The cells’ FUCCI signal is not shown, only H2B signal is shown faintly in the images, which instead show overlaid on the original image sequences the predominant (highest probability) fate predicted by the DEEP-MAP neural network through time at single-cell level for each cell detected. Magenta/blue/green/red: predicted hPSC/EC/ME/EN fate, correspondingly (colour code shown as image inset). EN triggered cells lose predicted hPSC status after just ∼1 day following differentiation trigger and evolve gradually toward the EN fate apparently transiting through an intermediate ME fate.

---

**Supplementary Figure 1.**
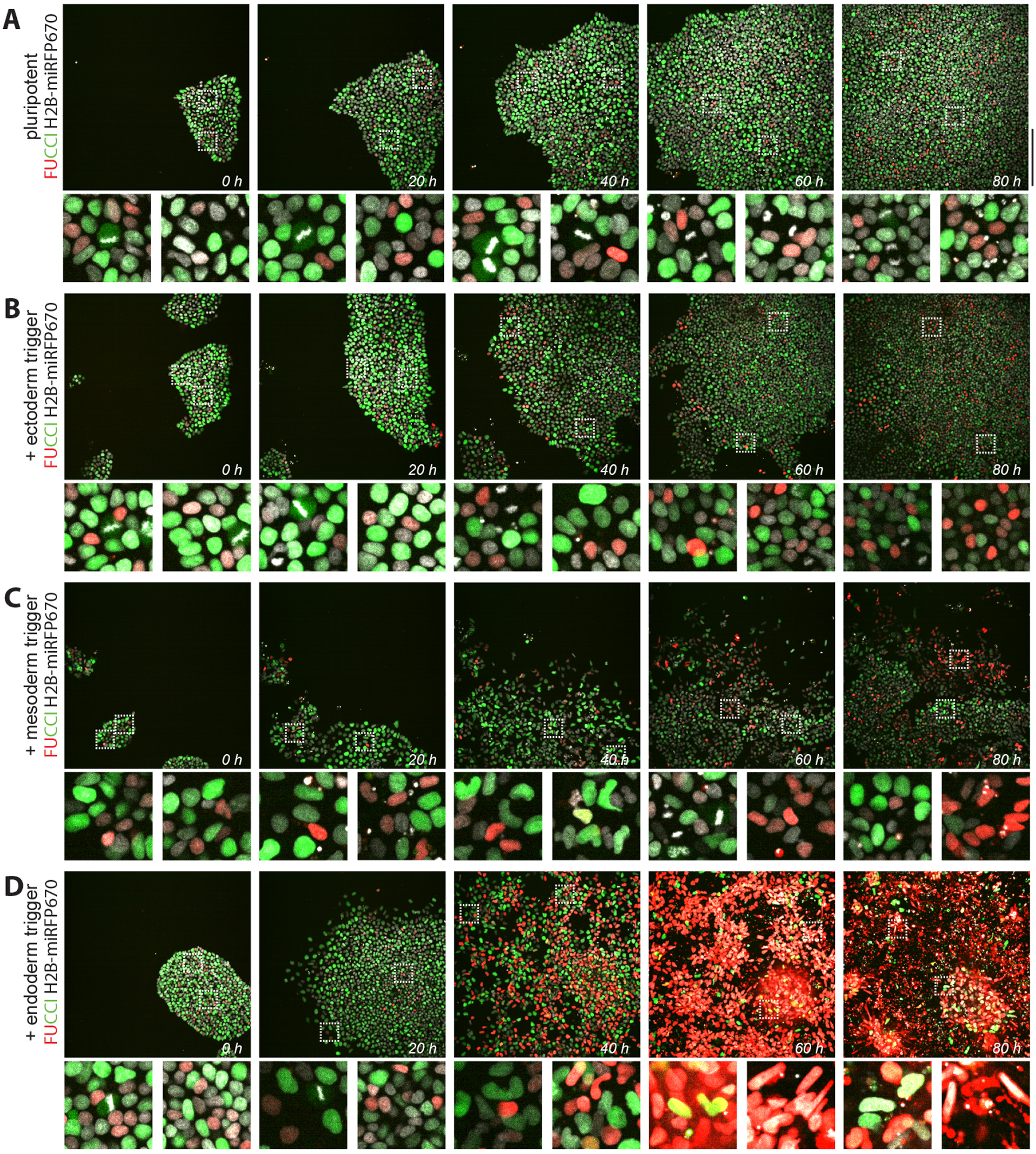
Changes in hPSC proliferation, morphology and cell cycle status during early differentiation. Image galleries of cells co-expressing FUCCI and the live chromatin reporter H2B-miRFP670 imaged continually by optimised, multi-day time-lapse microscopy for 5 days as they either maintain pluripotency (**A**) or after receiving trigger for ectoderm (**B**), mesoderm (**C**) or endoderm (**D**) differentiation at day 0. Dotted boxes in the top rows indicate areas magnified below. Images shown only from the first 80 hours of time-lapse. Scalebars: 100 μm. Note changes in the cells’ appearance after 24h of imaging among the different conditions, particularly ME and EN triggered cells that visibly become more spread out and alter their nuclear shape and cell cycle reporter characteristics.

**Supplemental Figure 2.**
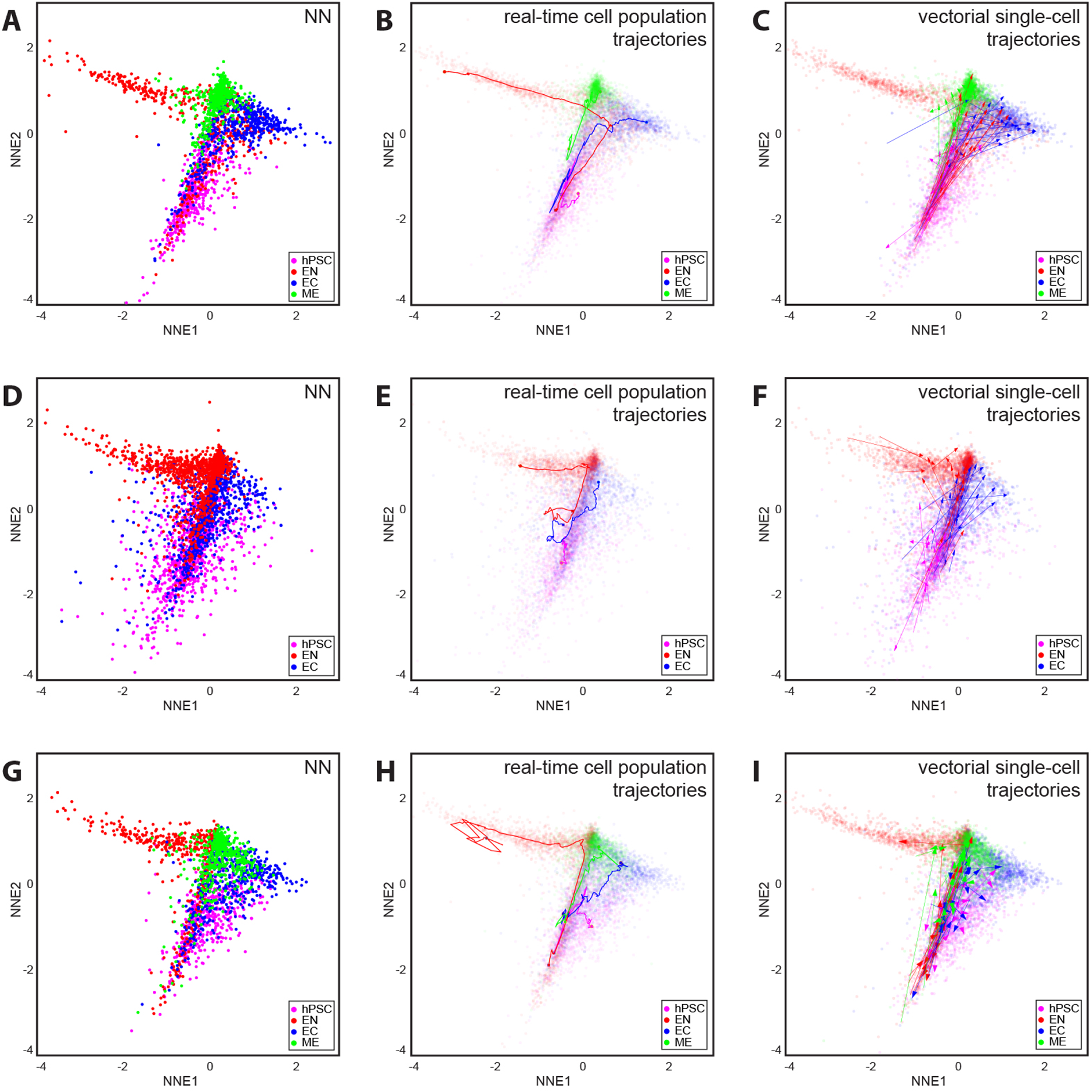
Neural networks allow reproducible mapping of cell fate dynamics toward multiple fate outcomes. A, D and G, Morphological phenoprints from three independent biological experiments showing neural networks reproducibly allow to clearly map and separate cell populations based on their proliferative phenoprints. All three mappings use the same neural network embedding model, which was trained with a subsample of data from (A). B, E and H, Maps showing the dynamical evolution and phenotypic diversity of the different cell fate populations in real-time, across the three independent experiments. Solid lines: population trajectories; magenta/red/blue/green: hPSC/EN/EC/ME cell populations, correspondingly. Real-time trajectories are shown as solid lines overlaid on the neural network embedding in E (made partly transparent for visualisation purposes). C, F and I, Vectorial trajectories of single-cells tracked early in differentiation along the different lineages, for the three independent experiments. Arrows: Vectorial trajectories; magenta/red/blue/green: hPSC/EN/EC/ME cell populations, correspondingly. Single-cell trajectories are shown as solid arrows overlaid on the neural network embeddings in A, D and G correspondingly, which are made partly transparent for visualisation purposes.

**Supplemental Figure 3.**
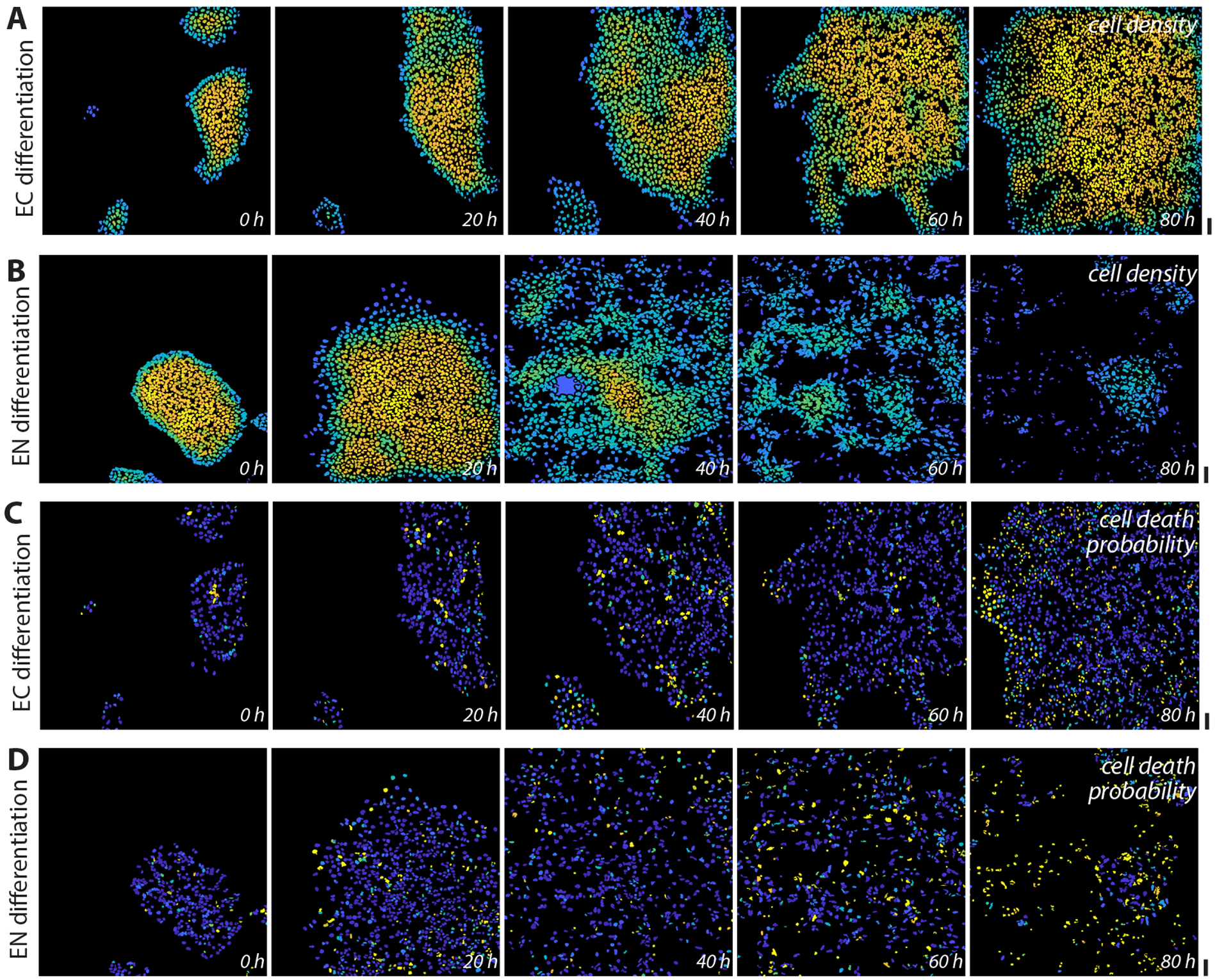
Visualising morphological and proliferative feature changes linked to cell fate changes. Image galleries of FUCCI and H2B-miRFP670 co-expressing cells imaged continually by optimised, multi-day time-lapse microscopy for 5 days after receiving trigger for ectoderm (**A, C**) or endoderm (**B, D**) differentiation at day 0. Image gallery corresponds to the same cells and colonies shown in Figure S1. Images shown only from the first 80 hours of time-lapse. FUCCI and H2B signals are not shown, instead images are fake coloured to display the feature value levels for cell density (**A, B**) or cell death probability (**C, D**) - two features shown in Figure 3 to have high importance in distinguishing different cell fates - through time at single-cell level for each cell detected. Note that EC cells show high density throughout (**A**), while in EN cells density changes dramatically (**B**). By contrast cell death probability increases in both EC (**C**) and EN (**D**) cells through time. Scalebars: 50 μm.

